# Yellow-Fruited Phenotype is Caused by an Insertion Event at 5’ UTR of *YFT1 Allele* in *yft1* Mutant Tomato ^1^

**DOI:** 10.1101/2020.05.04.077396

**Authors:** Weihua Zhao, Lei Gao, Yuhang Li, Minghui Wang, Lida Zhang, Lingxia Zhao

**Author notes:** Corresponding author: Lingxia Zhao, e-mail address, Department of Plant Science, School of Agriculture and Biology, Shanghai Jiao Tong University, 800 Dongchuan Road, Shanghai 200240, China. National Research and Development Center for Edible Fungus Processing Technology, Henan University, Jinming North Avenue, Kaifeng 475004, China (Lei Gao). Author Contributions L.Z. conceived and designed the research plans, and wrote the manuscript; W.Z. performed functional analysis of *YFT1* through genetic transformation, chromoplast development using TEM, and promoter strength analysis, and wrote the manuscript; Y.L. examined carotenoids accumulation using UPC^2^; L.G., M.W. and L.Z. participated in transcriptional analysis.

## Abstract

The *yft1* tomato mutant has a yellow-fruited phenotype controlled by a recessive gene (*YFT1*), which has been shown by map-based cloning to be a homolog of *ETHYLENE INSENSITIVE 2* (*EIN2*). The genetic lesion of the *YFT1* allele of *yft1* is attributed to a 573 bp DNA fragment (IF_573_) insertion at 1,200 bp downstream of the transcription start site (TSS). Transcriptome analysis revealed that the mutation resulted in 5,053 differentially expressed genes (DEGs) in the fruit pericarp compared with the M82 wild type cultivar. These were annotated as being involved in ethylene synthesis, chromoplast development, and carotenoid synthesis. Genetic lesion in *YFT1* caused a reduction in its own transcript levels in *yft1* and impaired ethylene emission and signal transduction, delayed chromoplast development and decreased carotenoid accumulation. The molecular mechanism underlying the reduced expression of *YFT1* in *yft1* was examined at both the RNA and DNA levels. The IF_573_ event was shown to introduce two negative regulatory sequences located at -272 to -173 bp and -172 to -73 bp in the *YFT1* allele promoter, causing alterative splicing due to aberrant splicing sites, and also altering the structure of the open reading frame in the 5’-UTR. This study contributes to the understanding of color formation in tomato fruit.

**One-sentence summary:** Lesion happened in regulatory region impairs expression of a key gene of ethylene signal pathway, and alters fruit color in tomato due to effect of carotenoids accumulation and ethylene synthesis.

## Introduction

The gaseous plant hormone ethylene has a range of effects on diverse physiological and developmental processes, including the induction of the seedling triple response, the promotion of seed germination, cell division, leaf epinasty, leaf and flower senescence and abscission and climacteric fruit ripening, as well as the inhibition of cell elongation (Ecker, 1995; Giovannoni, 2001; Ju et al., 2015), particularly affects the ripening and coloration of fruit (Lanahan et al., 1994; Vrebalov et al., 2002; Karlova et al., 2014; Zhu et al., 2014). Ethylene also acts as a signal molecule in response to pathogen challenge and wounding (Foroud et al., 2019; Tsolakidou et al., 2019).

A complex ethylene signal transduction pathway has been identified, involving ethylene binding by, and inactivation of, the endoplasmic reticulum (ER)-localized ETHYLENE RECEPTOR (ETR) proteins, thereby affecting their interaction the dominant negative regulator protein kinase CONSTITUTIVE TRIPLE RESPONSE 1 (CTR1) (Hua et al., 1995; Qiao et al., 2012). CTR1 phosphorylates ETHYLENE INSENSITIVE 2 (EIN2), leading to its degradation *via* the P26S protein complex. In the presence of ethylene, EIN2 is dephosphorylated at Ser^645^ and Ser^924^ and the cleaved C-terminal (CEND) proteolytic fragment traffics from the ER to the nucleus to form a complex with EIN3/EIL (ETHYLENE INSENSITIVE 3 LIKE). This complex binds to ETHYLENE RESPONSIVE ELEMENTS (EREs) in the promoters of target genes to regulate their expression (Ju et al., 2012; Qiao et al., 2012). Li et al. (2015) found that partial CENDs retained in the cytoplasm can form a complex with EIN5 and PABs (poly A binding proteins), which regulates the expression of *EIN3-BINDING F-BOX 1*(*EBF1*)/*EBF2* at the translational level via binding to its 3’ UTR region. Thus, EIN2 acts as a core component in ethylene signal transduction.

An important experimental system for studying ethylene regulation has been tomato (*Solanum lycopersicum*) fruit, which exhibit a major increase in ethylene emission during ripening, in parallel with changes in color, aroma, texture and profiles of primary and secondary metabolites (Karlov et al., 2014; Liu et al., 2015). Mechanistic insights into ripening regulation have resulted from the characterization of tomato fruit ripening mutants *ripening inhibitor* (*rin*) and *non-ripening* (*nor*), which are perturbed in multiple aspects of ripening, including ethylene biosynthesis and production of the carotenoid pigment lycopene (Vrebalov et al., 2002; Manning et al., 2006). In addition, the tomato ethylene receptor mutant *Nr* (*Never ripe*) is insensitive to ethylene and has fruit that fail to fully ripen (Wilkinson et al., 1995; Hackett et al., 2000), and that accumulate substantial reduced levels of lycopene and its derivative β-carotene (Lanahan et al., 1994). The generation and analysis of transgenic tomato plants has also provided insights into ethylene signaling in fruit ripening, such as the demonstration that down-regulation of *LeETR4* accelerates ripening (Tieman et al., 2000).

Tomato is an economically important crop, with an annual production of 182 million tons and a value of $80 billion in 2017 (http://www.fao.org/faostat/en/#data/QV/visualize). The fruit contain many nutritionally important compounds, including beta-carotene, which is the major precursor for vitamin A synthesis, (DellaPenna and Pogson, 2006), and the antioxidant lycopene, which can be help reduce the incidence of chronic diseases (Ford and Erdman, 2012). Fruit color (predominantly red, pink and yellow) is determined by the content of carotenoids, especially the ratio of lycopene to β-carotene (Isaacson et al., 2002; Enfissi et al., 2017; Chen et al., 2019), and has been associated with fruit nutritional value as well as appearance. The carotenoid synthesis pathway has been well characterized at both biochemical and molecular levels (Fray and Grierson, 1993; Ronen et al., 2000; Hirschberg, 2001; Isaacson et al., 2002; Galpaz et al., 2008; Kachanovsky et al., 2012; Neuman et al., 2014; Pankratov et al., 2016); however, the mechanisms of its regulation are less understood.

A *yellow fruited tomato 1* (*yft1*) mutant (also named *n3122*) was shown, by map based cloning, to have a genetic lesion comprising a 573 bp insertion and a 13 bp deletion in the promoter region (−318 bp upstream of the ATG start codon) of an *EIN2* gene (Gao et al., 2016). Expression of *YFT1* allele in *yft1* was significantly down-regulated compared to the wild type cultivar M82 and resulted in a yellow-fruited phenotype (Gao et al., 2016). However, the precise genetic evidence that *YFT1* controls fruit color formation in tomato is still lacking, and the molecular mechanism behind its downregulation in *yft1* remains to be determined.

To this end, here we performed *YFT1* functional complementation, large-scale transcriptomic analysis and functional dissection of the mutated control regions/promoter of the *YFT1* allele to investigate the involvement of *YFT1* in tomato fruit color formation. Our work provides new insights into the regulation of tomato fruit color formation, and particularly the interaction between carotenoid and ethylene synthesis and signal transduction.

## Results

### Genetic lesion of *YFT1* in *yft1* causes a yellow-fruited phenotype due to an insertion of 573 bp DNA

The yellow-fruited phenotype of *yft1* was previously inferred to be due to a genetic lesion in a homolog of *EIN2* (*Solyc09g007870*) (Gao et al., 2016), but there still is lack of genetic evidences, and the underling mechanism of the down-regulated expression of the *YFT1* allele in *yft1* mutant had remained elusive yet.

*YFT1* is preferentially expressed in reproductive organs in the M82 cultivar, especially in the stamen and pistil. However, in the *yft1* mutant, *YFT1* was observed to be preferentially expressed in vegetative organs, with expression being particularly reduced in stamens and fruit (Supplemental Figure S1). *YFT1* expression was also found to increase with fruit development in M82, where it was significantly higher than in *yft1* fruit (Supplemental Figure S1).

Constitutive overexpression of *YFT1* in the *yft1* mutant, by introduction of the *YFT1-CDS* construct, rescued the yellow-fruited phenotype, whereas silencing *YFT1* expression in M82 by RNA-interference (RNAi) using *YFT1-RNAi* resulted in a yellow-fruited phenotype and delayed fruit development, similar to the phenotypes seen in *yft1* (Fig. 1). *YFT1* expression was significantly higher in the *YFT1-CDS* lines than in *yft1*, and even higher than in M82. The expression level of *YFT1* in the *YFT1-RNAi* lines was statistically lower than in M82, whereas it was significantly higher than in *yft1* (Fig. 2). *YFT1* expression also increased with fruit development in both *YFT1-CDS* and M82, and was significantly higher than in *yft1* and *YFT1-RNAi* (Fig. 2). These results suggested that overexpression of *YFT1-CDS* in *yft1* was sufficient to generate a typical a red-fruited phenotype.

**Figure 1.**
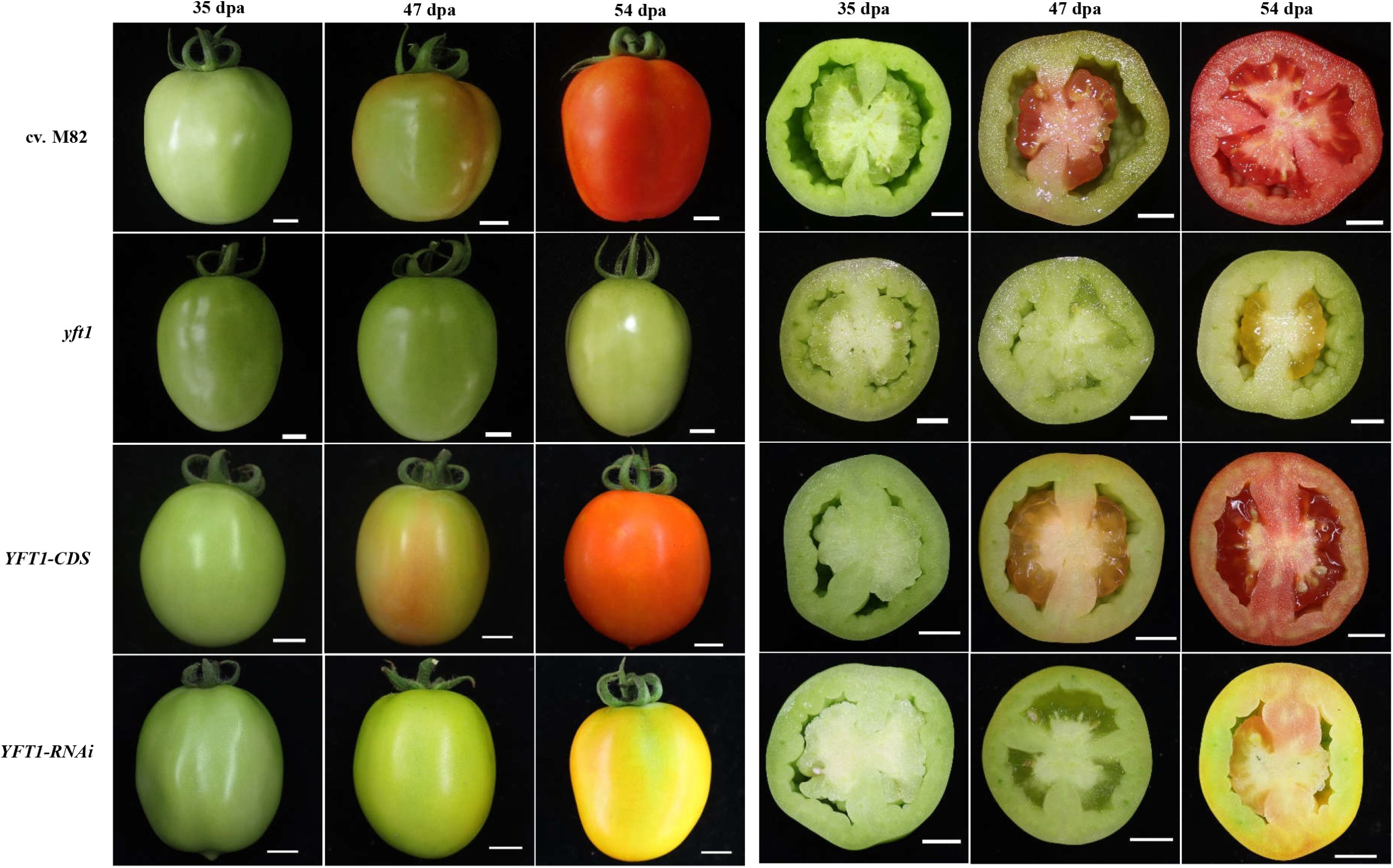
Fruit color of the tomato lines with different genetic background during developmental stages. **cv. M82**, a cultivated tomato (*S. lycopersicum*), which serves as wild type (WT) in present study. ***yft1***, yellow-fruited tomato 1 (n3122), which created from cv. M82 via fast-neutron irradiation. ***YFT1-CDS***, overexpressed *YFT1* in *yft1* via genetic transformed using construct of *2*×*35S::YFT1-CDS*. ***YFT1-RNAi***, knockdown expression of *YFT1* in cv.M82 via genetic transformed using construct of *35S::YFT1-RNAi* by RNA interference technique. Bars=1 cm.

**Figure 2.**
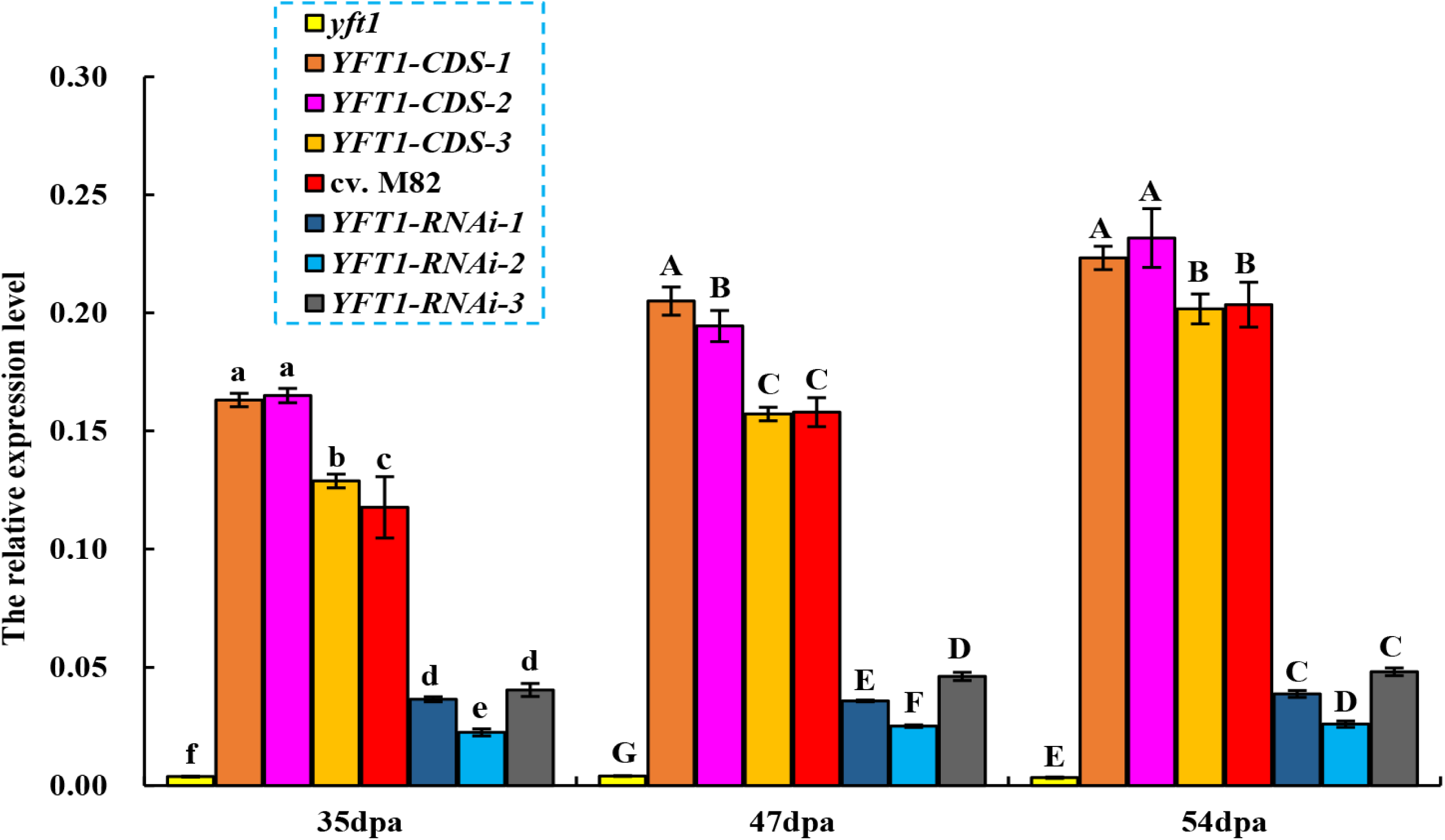
*YFT1* expression in the *yft1, YFT1-CDS*, M82 and *YFT1-RNAi* lines. **M82**, a cultivated tomato (*S. lycopersicum*) cultivar, served as wild type (WT). ***yft1***, *yellow-fruited tomato 1* (*n3122*), a mutant created from M82 via fast-neutron irradiation. ***YFT1-CDS***, overexpressing *YFT1* coding sequence (CDS) in *yft1* via genetic transformation. ***YFT1-CDS* (1-3)**, different *YFT1-CDS* lines. ***YFT1-RNAi***, knockdown of *YFT1* expression in M82 *via* RNA interference. ***YFT1-RNAi* (1-3)**, different *YFT1-RNAi* lines. Data values indicate the mean ±SD (n=3). Capital and lower-case letters indicate statistical signifcance at *P*<0.01 and *P*<0.05 determined by a Duncan’s test.

The genetic lesion in the *YFT1* allele on chromosome 9 in *yft1* comprises two In/Del events: an insertion of 573 bp and a deletion of 13 bp. The 573 bp insertion segment (referred to as *IF*_*573*_) sequence was found to be identical to a DNA sequence on chromosome 6 (https://www.solgenomics.net/) located in a 9,845 bp region between *Nodulin-like* (*Solyc06g082410.1*) and *Peroxidase 3* (*Solyc06g082420.2*). There was, however, no mutation or loss of nucleotides in *yft1* compared with M82 in this region of chromosome 6 when it was amplified and sequenced using specific primers (Ch06-573u-F/Ch06-573d-R) (Supplemental Figure S2 and Supplemental Table S1). This indicated that the yellow-fruited phenotype was caused by the genetic lesion in *YFT1* on chromosome 9, and not by loss of DNA segment or mutation on chromosome 6 (Supplemental Figure S2).

To characterize the effect of both *IF*_*573*_ and the 13 bp deletion on *YFT1* expression, a series of artificial promoters (*pYFT1, pyft1, pYFT1-Del13, pyft1*+*Del13u*, and *pyft1*+*Del13d*) were created, and fused to upstream of the *GUS* gene. These constructs were transiently expressed in tobacco, and while a difference in GUS activity was not detected between *pYFT1* and *pYFT1-Del13*, or *pyft1, pyft1*+*Del13u* and *pyft1*+*Del13d*. However, the GUS activity driven by *pYFT1* or *pYFT1-Del13* was significantly higher than for *pyft1, pyft1*+*Del13u* or *pyft1*+*Del13d.* This suggested that the downregulation of *YFT1* in *yft1* could be primarily attributed to the 573 bp insertion in the promoter region, and not the 13 bp deletion (Supplemental Figure S3).

### Genetic lesion of the *YFT1* allele in *yft1* has an effect on transcriptomic profiling

Transcriptomic analyses of pericarp samples from *yft1* and M82 were collected at 35 (Mature green), 47 (Breaker) and 54 (Red ripe) dpa revealed that the genetic lesion of *YFT1* resulted in total of 5,053 DEGs in *yft1* compared to M82. These comprised 329, 3610 and 3751 DEGs at 35, 47 and 54 dpa, respectively (Fig. 3A). Pathway enrichment analysis suggested that the lesion in *YFT1* affected numerous physiological, biochemical and developmental processes. These included ethylene signal transduction, and the metabolism of cysteine and methionine, which are used for ethylene and carotenoid synthesis (Fig. 3B). These results are consistent with the *YFT1* lesion affecting fruit color in *yft1* through the regulation of ethylene signal transduction and carotenoid synthesis (Fig. 3C).

**Figure 3.**
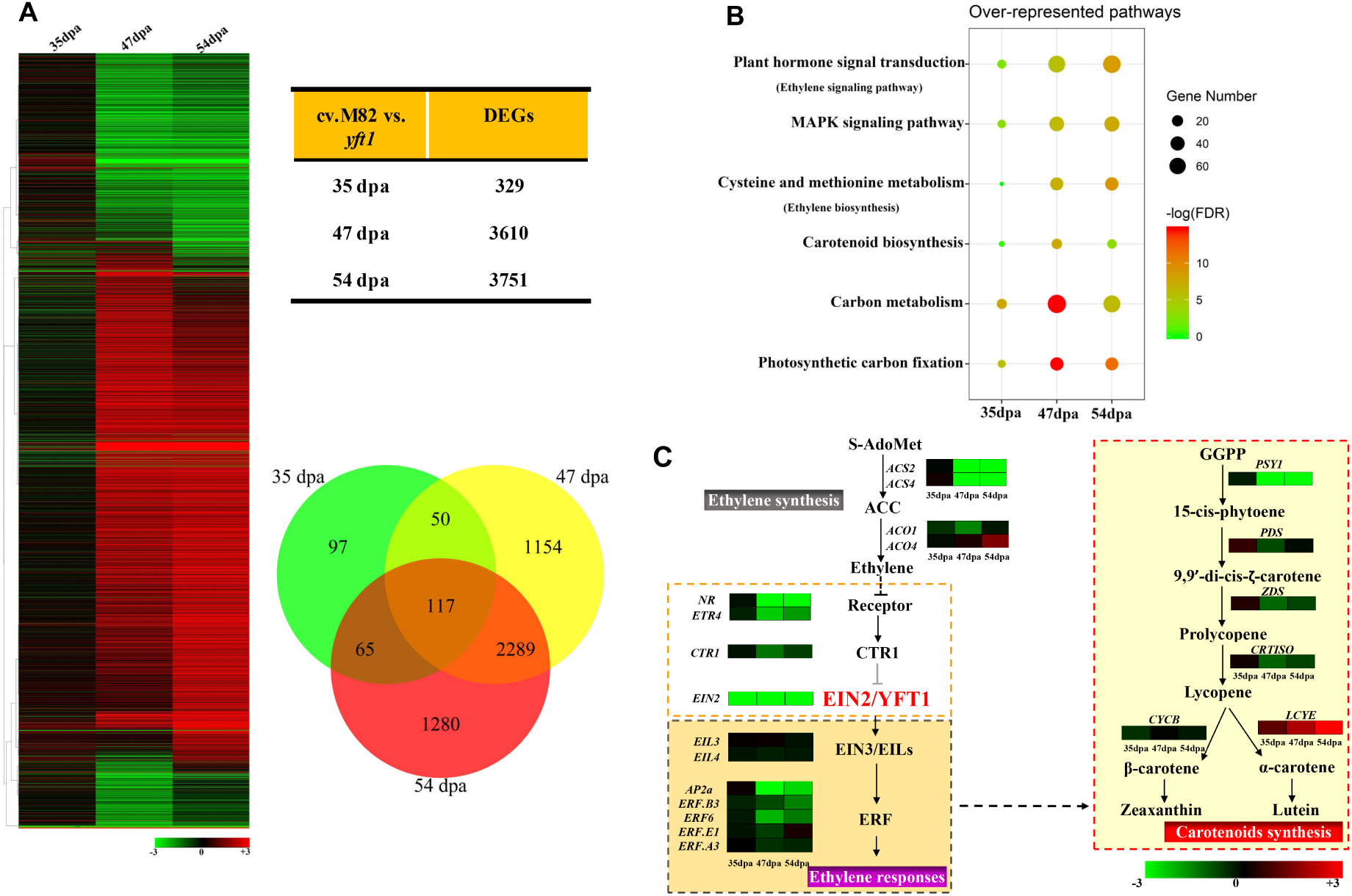
Transcriptome analysis pericarp from *yft1* and M82 at different fruit development stages. **(A)** Cluster of differentially expressed genes (DEGs) between M82 and *yft1* during fruit development. (**B**) Genetic lesion of *YFT1* affects main biological pathways in *yft1* compared to M82. **(C)** Key genes involved in ethylene biosynthesis and signal transduction, and carotenoid biosynthesis. **DEG**s, differentially expressed genes. **FDR**, false discovery rate. **ACC**, 1-aminocyclopropane-1carboxylic acid. **ACO**, ACC oxidase. **ACS**, ACC synthase. **CTR**, constitutive triple-response. **EIN2**, Ethylene insensitive 2. **EIN3/EIL**, ethylene insensitive 3-like. **ETR**, ethylene receptors. **ERF**, ethylene response factor. **GGPP**, geranylgeranyl pyrophosphat. **NR**, Never ripe. **S-adoMet**, S-adenosylmethionine.

### *YFT1* affects chromoplast development and carotenoid accumulation

Chromoplasts are a specialized type of plastids in which carotenoids accumulate and that develop in flowers and fruits, thereby contributing to their color. We previously reported that chromoplast development in *yft1* is delayed (Gao et al., 2016). When we used transmission electron microscopy (TEM) to examine chromoplast development in the various transgenic lines developed through this current study, we observed that the chloroplast envelope and grana (gr) edges were clearly distinguishable in fruit at 35 dpa, and that there was no differences in chromoplast structure between the four *YFT1-CDS, YFT1-RNAi*, M82, and *yft1* lines (Fig. 4). We observed that during fruit development, the chloroplast envelope membrane and thylakoid structure degraded, and at 47 dpa the linear thylakoid membrane(lth) was intact in both M82 and *YFT1-CDS*, but was completely gone at 54 dpa (RR stage), and even formed *carotenoid crystalloid structures* (ccr) accumulated more carotenoids. In contrast, the chloroplast envelope membrane and *grana* (gr) structures in both *yft1* and *YFT1-RNAi* were easily distinguishable at 47 dpa and 54 dpa (Fig. 4).

The number of plastoglobules (pg) containing carotenoids increased during fruit development and while few pg were detected at 35 dpa in all four tomato lines, there were more in both M82 and *YFT1-CDS* than in either *yft1* and *YFT1-RNAi* (Fig. 4). We also observed that the number of pgs based on the TEM images correlated with the levels of carotenoids in the pericarp, and peaked at 54 dpa (Fig. 4). The number of pgs in both M82 and *YFT1-CDS* was considerably higher than in *yft1* and *YFT1*-*RNAi* (Fig. 4), and ccrs were also observed in both M82 and *YFT1-CDS* (Fig. 4).

**Figure 4.**
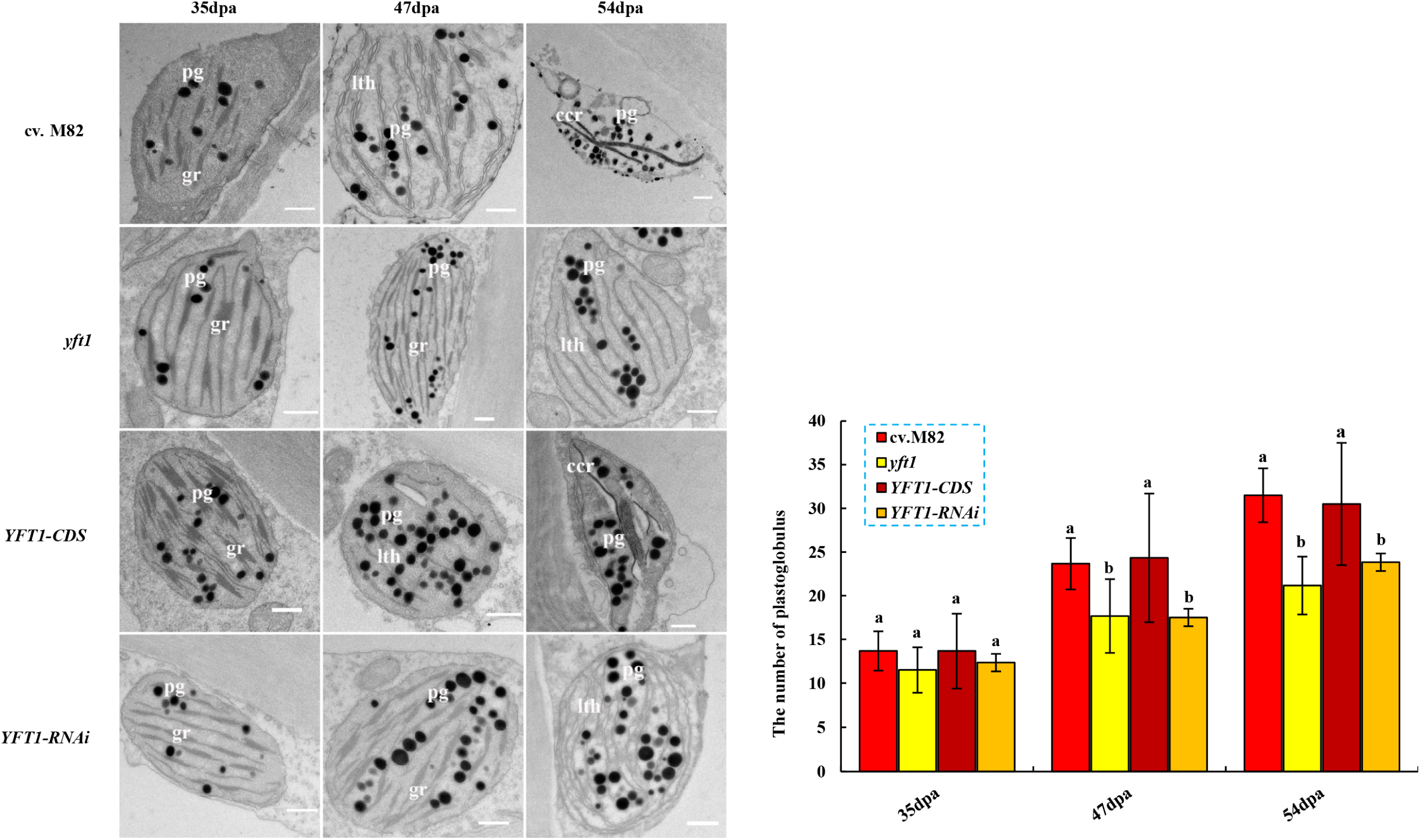
The difference of plastoglobule formation and number through chromoplast developmental visualized by transition electron microscopy (TEM) **M82**, wild type tomato. ***yft1***, *yellow-fruited tomato 1* (*n3122*), a mutant created from M82 via fast-neutron irradiation. ***YFT1-CDS***, overexpressing *YFT1* in *yft1*. ***YFT1-RNAi***, knockdown of *YFT1* expression in M82 by RNA interference. ***pg***, plastoglobule. ***gr***, grana. ***lth***, long linear thylakoid membrane structure. ***ccr***, carotenoid crystalloid structure. Scale bars = 500 nm. Data are the means ± SD (n =3). Lowercase letters indicate statistical signifcance at *P*<0.05 as determined by a Duncan’s test.

The effect of the *YFT1* genetic lesion on carotenoid accumulation was also examined using Ultra-performance Convergence Chromatography (UPC^2^). The lycopene content increased with fruit development, and peaked at 54 dpa (RR) in all four tomato lines (Fig. 5A). However, lycopene accumulation in both M82 and *YFT1-CDS* was significantly higher than in *yft1* or *YFT1-RNAi* at 54 dpa, and little or no lycopene was detected at both 35 dpa (MG) and 47 dpa (BR) (Fig. 5A). The β-carotene content also increased with fruit development and was higher in both M82 and *YFT1-CDS* than in *yft1* or *YFT1-RNAi* at both 47 dpa and 54 dpa, but not at 35 dpa (Fig. 5A). In contrast, levels of both α-carotene and lutein reached a peak at 47 dpa and then decreased. Notably, α-carotene was not detected in any of the four tomato lines at 35 dpa, and significantly higher levels were present in M82 and *YFT1-CDS* than in *yft1* and *YFT1-RNAi* at both 47 dpa and 54 dpa (Fig. 5A). The concentration of lutein was significantly higher in M82 and *YFT1-CDS* than in *yft1* or *YFT1-RNAi* fruit (Fig. 5A).

**Figure 5.**
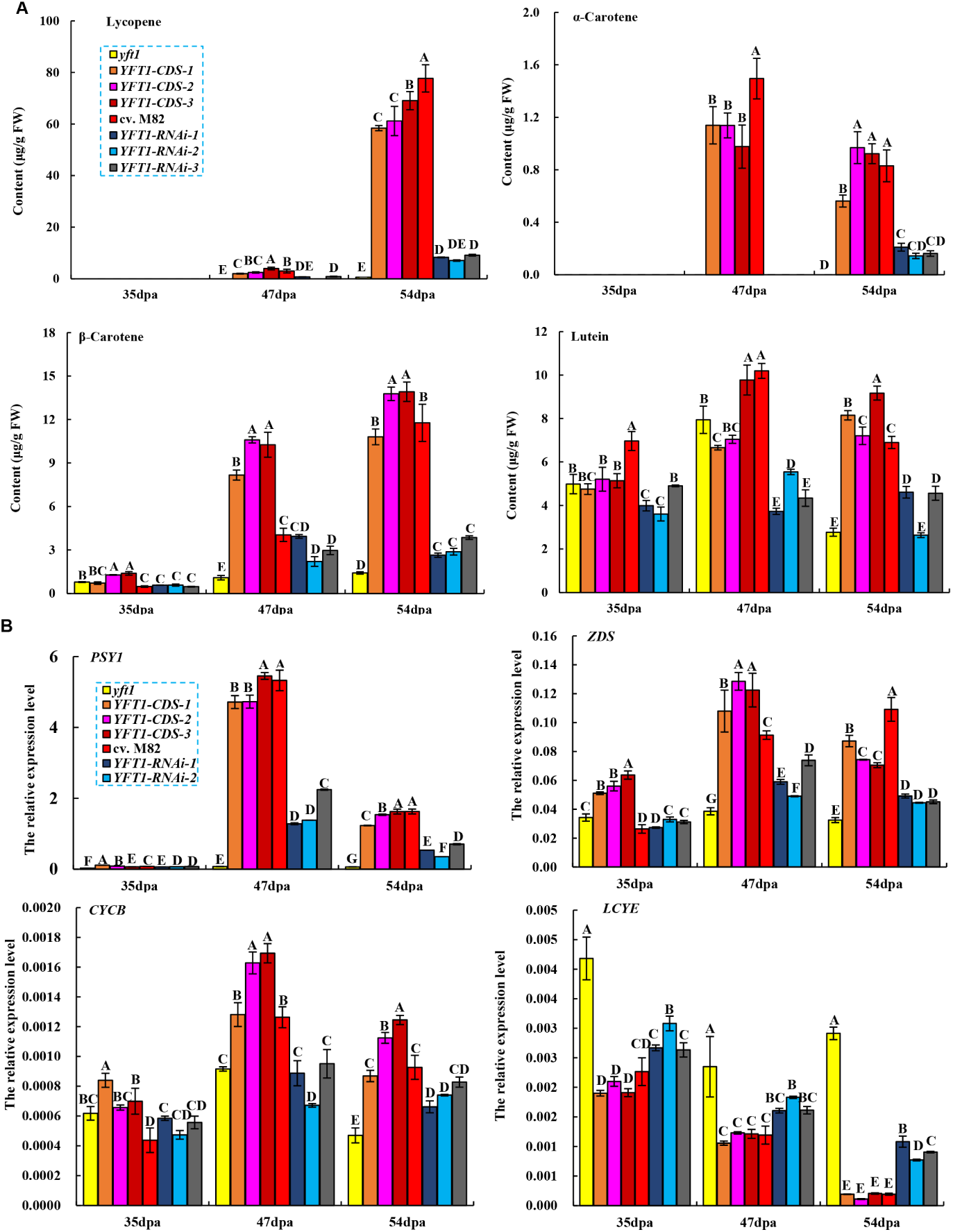
The *YFT1* genetic lesion decreased carotenoid accumulation. **(A)** Carotenoid accumulation in tomato lines at different developmental stages. **(B)** Expression of the *PSY1, ZDS, LCYE* and *CYCB* genes, involved in the carotenoid synthesis pathway. **M82**, wild type tomato. ***yft1***, *yellow-fruited tomato 1* (*n3122*), a mutant created from M82 via fast-neutron irradiation. ***YFT1-CDS***, overexpressing *YFT1* in *yft1*. ***YFT1-RNAi***, knockdown of *YFT1* expression in M82 by RNA interference. ***PSY1***, gene encoding phytoene synthase 1 protein. ***ZDS***, gene encoding ξ-carotene desaturase. ***LCYE***, gene encoding lycopene ε-cyclase. ***CYCB***, gene encoding chromoplast specific lycopene β-cyclase. Data are the means ± SD (n =3). Capital letters indicate statistical signifcance at *P*<0.01 as determined by a Duncan’s test.

We next measured the expression of genes in the carotenoid synthesis pathway. Both *PSY1* (*phytoene synthase 1*) and *ZDS* (*ξ-carotene desaturase*) encode enzymes that catalyze the synthesis of linear carotenoids, while *LCYE* (*lycopene ε-cyclase*) and *CYCB* (*chromoplast specific lycopene β-cyclase*) are involved in the production of cyclic carotenoids. *LCYE* expression decreased during fruit development, while expression levels of *PSY1, ZDS*, and *CYCB* reached a peak at 47 dpa, and were significantly lower at both 35 dpa and 54 dpa (Fig. 5B). The expression of *PSY1* was significantly higher in both M82 and *YFT1-CDS* than in *YFT1-RNAi* or *yft1*, especially at 47 dpa (BR) where it reached a peak, whereas expression was low or undetectable at 35 dpa. In contrast, *PSY1* expression was not detected in *yft1* (Fig. 5B). *ZDS* and *CYCB* were also expressed in significantly higher levels in both M82 and *YFT1-CDS* than in *yft1* or *YFT-RNAi* at 47 dpa and 54 dpa (Fig. 5B). *LCYE* expression was significantly higher in *yft1* and *YFT1-RNAi* than in either M82 or *YFT1-CDS* (Fig. 5B). This suggested that the genetic lesion in *YFT1* resulted in a reduction in the expression of genes involved in linear carotenoid synthesis but an increase in the expression of those involved in the synthesis of cyclic carotenoids.

### The genetic lesion in *YFT1* alters the biological behavior of ethylene in *yft1*

To examine the effects of ethylene on *yft1* seedling development, we germinated seedlings on MS_0_ medium containing the ethylene precursor 1-aminocyclopropane-1-carboxylic acid (ACC). Seedling phenotypes were examined 7 days after sowing. In the absence of ACC, seedlings from all four tomato lines were hookless, grew rapidly and had thin stems and roots, and numerous lateral roots (Fig. 6A). In contrast, in the presence of 10 μM ACC, seedlings had apical hooks, stem and root growth was slow, and there were no lateral roots or root hairs. We also observed that the *YFT1-CDS* seedlings were significantly smaller than M82, *yft1* and *YFT1-RNAi* seedlings, regardless of whether supplemental ACC was present or not (Fig. 6A), possibly due to the constitutive promoter of the *2*×*35S::YFT1-CDS* construct mediating strong ethylene signal transduction.

**Figure 6.**
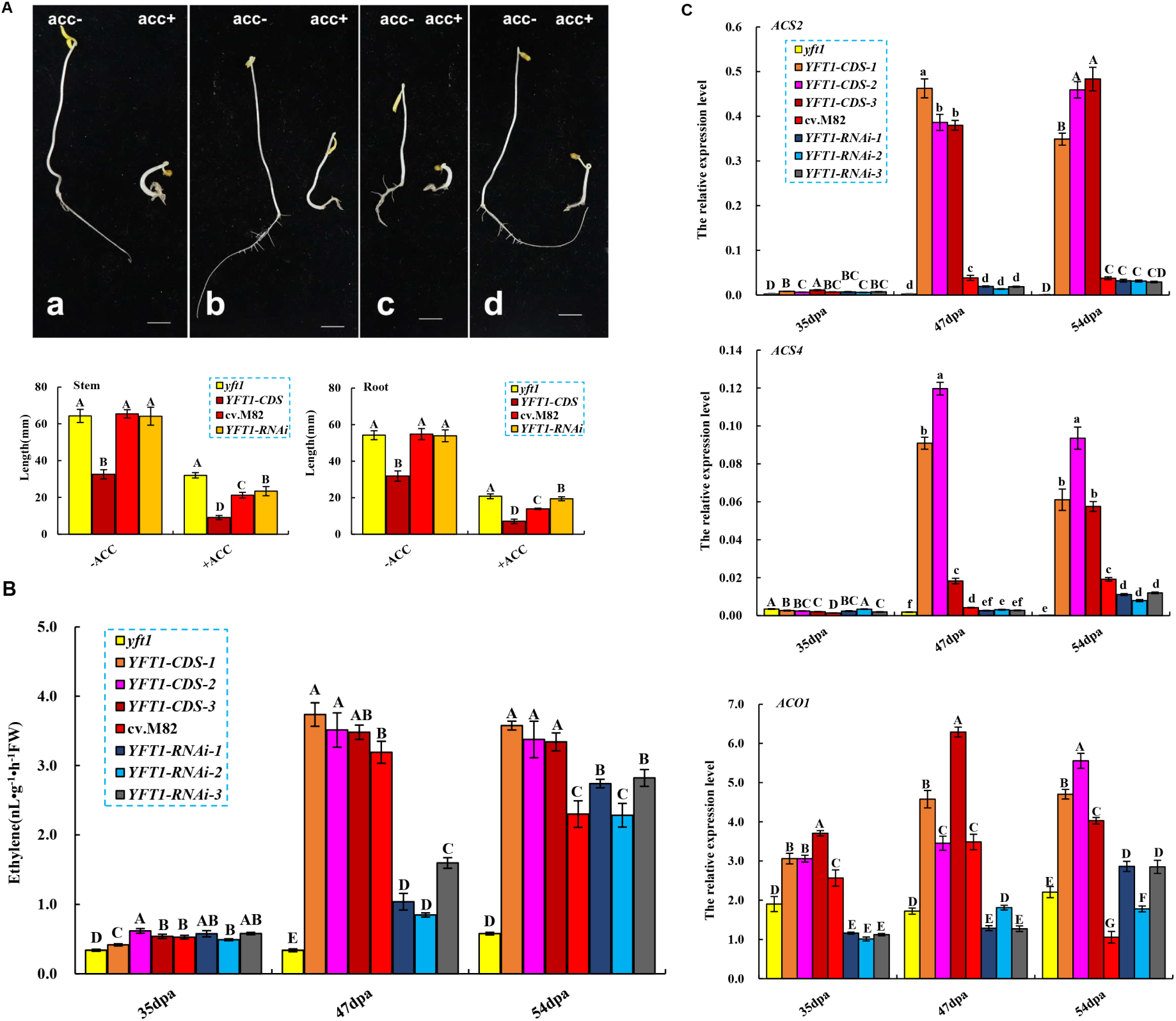
The regulation of *YFT1* on ethylene emission and signaling transduction. **(A)** The seedling phenotypes after 7 days on MS_0_ medium with or without ACC (10 μM). (a) M82. (b) *yft1*. (c) *YFT1-CDS*. (d) *YFT1-RNAi*. **(B)** Difference in ethylene emission between the tomato lines with different genetic backgrounds at different developmental stages. **(C)** Expression of *ACO1* and *ACS2/4* in tomato lines with different genetic backgrounds. **M82**, wild type tomato. ***yft1***, *yellow-fruited tomato 1* (*n3122*), a mutant created from M82 via fast-neutron irradiation. ***YFT1-CDS***, overexpression of *YFT1* in *yft1*. ***YFT1-RNAi***, knockdown of *YFT* expression in M82 using RNAi. Data indicate the means ± SD (n =3). Capital and lower-case letters indicate statistical signifcance at *P*<0.01 and *P*<0.05 as determined by a Duncan’s test.

In *yft1*, ethylene emission was constant throughout fruit development and at relative low levels, while it increased in the other lines. A climacteric peak was observed at 47 dpa (BR stage), that was much higher in M82 and *YFT1-CDS* than in *YFT1-RNAi* and *yft1* (Fig. 6B). To further investigate the relationship between the genetic lesion of *YFT1* in *yft1* and ethylene synthesis we measured the expression of genes encoding the ethylene biosynthetic enzymes ACC synthase (ACS) and ACC oxidase (ACO). *ACS2/4* were exclusively expressed in ripening *YFT1-CDS* fruit at 47 dpa (BR) and 54 dpa (RR), but not in green mature fruits (35 dpa), and expression was also higher than in the other three lines, despite the fact that expression differences were also found among the different *YFT1-CDS* lines (Fig. 6C). *ACS4* expression in M82 was significantly higher than in both *yft1* and *YFT1-RNAi*, but this was not the case for *ACS2* (Fig. 6C). *ACO1* expression was significantly higher in M82 and *YFT1-CDS* than in *yft1* and *YFT1-RNAi*, and reached a peak at 47 dpa in M82 and *YFT1-CDS*, whereas in *yft1* and *YFT1-RNAi* it increased steadily with fruit development (Fig. 6C).

### The reduced expression of *YFT1* expression in *yft1* involves two promoter regions

To dissect the functional regions of *IF*_*573*_, we generated a series of artificial promoter sequences containing different *IF*_*573*_ DNA segments fused to the *35S* promoter driving expression of the *LUCIFERASE* (*LUC*) gene. The strength of these artificial promoters was determined by transforming the constructs into tobacco plants and measuring LUC activity. We observed that *IF*_*573*_ did not drive *LUC* expression, and the insertion events between *35S* promoter and *LUC* would impair *LUC* expression (Fig. 7). LUC activities as a result of *35S-IF*_*73*_ and *35S-IF*_*173*_ was higher than those associated with the other four *35S-IF*_*273*_, *35S-IF*_*373*_, *35S-IF*_*473*_, and *35S-IF*_*573*_ promoters. However, the activity of *35S-IF*_*173*_ was higher than that of *35S-IF*_*73*_, suggesting that negatively regulatory sequences reside in the regions -272 to -173 bp and -172 to -73 bp regions of *IF*_*573*_ (Fig. 7).

**Figure 7.**
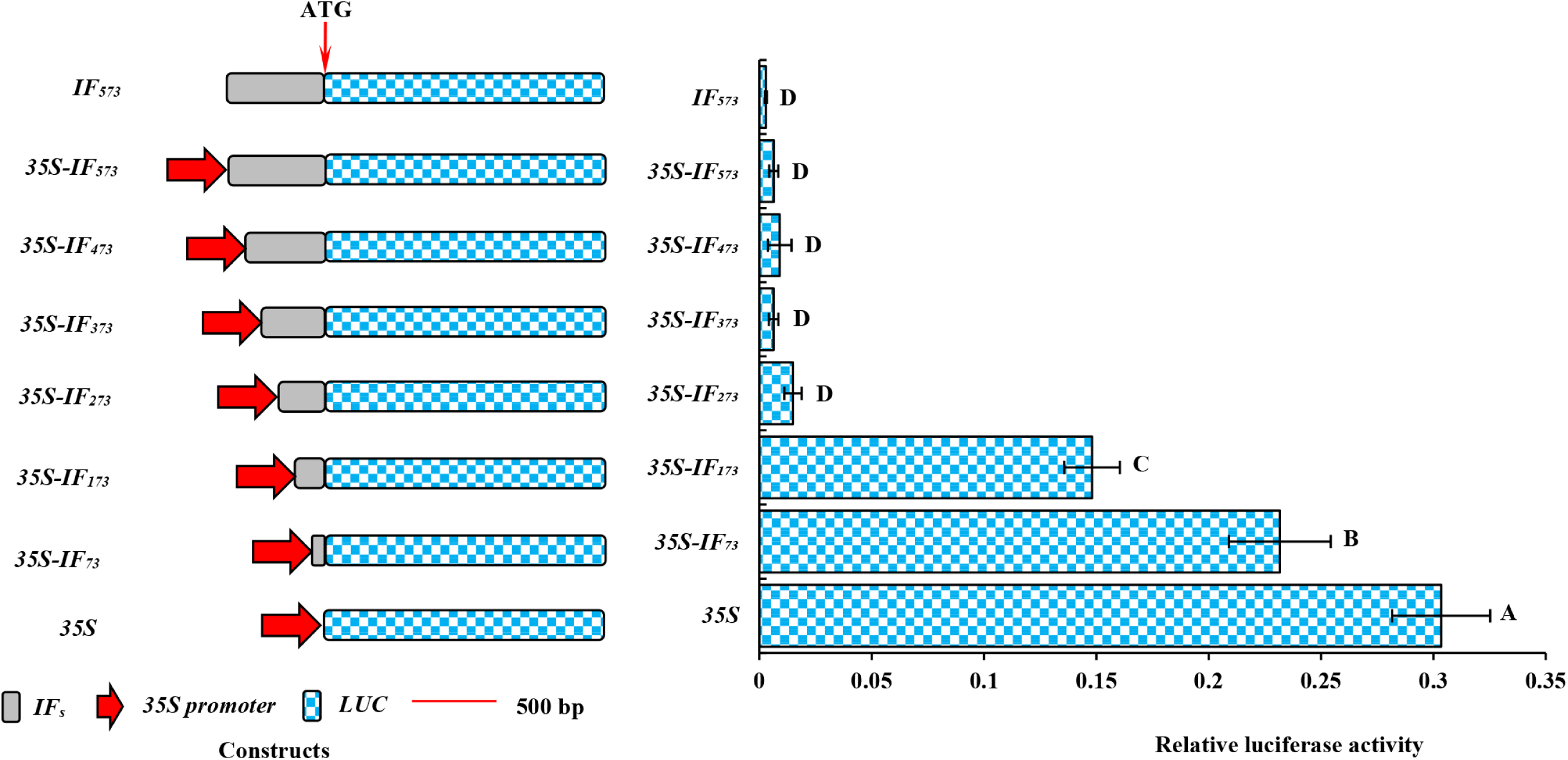
Dissection of the functional *IF*_*573*_ region. Constructs used (left) and luciferase activities (right). ***35S***, CaMV 35S promoter. ***IF***_***573***_, the 573 bp inserted DNA fragment. ***35S-IF***_***573***_, ***35S-IF***_***473***_, ***35S-IF***_***373***_, ***35S-IF***_***273***_, ***35S-IF***_***173***_ and ***35S-IF***_***73***_, indicate the gradual deletion of *IF*_*573*_ (100 bp each step) from left to right and downstream fusion to the *35S* promoter, and the subscript numbers following ‘*IF*’ in each construct indicate length of the remaining DNA from the right border of *IF*_*573*_. Data indicate the means ± SD (n=3). Capital letters indicate statistical significance at the *P* <0.01 level based on a Duncan’s test.

### The *IF*_*573*_ region causes aberrant splicing in the 5’-UTR

Transcription expression focused on the different regions in the *YFT1* allele of *yft1* appeared that expression level of upstream sequence of the inserted point (−318 bp upstream of ATG start codon) was dramatically higher than that in downstream, but not in M82 (Fig. 8A). This suggested that insertion of the 573 bp DNA segment causes alternative splicing of *YFT1* allele transcripts in *yft1*. To test this hypothesis, full length *YFT1* cDNA sequences were amplified from M82 and *yft1* using the RACE approach. A full length 4,850 bp cDNA was obtained from M82, consisting of a 647 bp 5′-UTR, a 3,951 bp CDS and a 260 bp 3′ UTR, which would encode a protein of 1,316 amino acids. In contrast, three different transcripts (*cDNAyft1.1, cDNAyft1.2* and *cDNAyft1. 3*) were amplified from *yft1*. The 458 bp *cDNAyft1.1* contained no CDS, suggesting that it does not encode a functional protein. Both *cDNAyft1.2* (5,418 bp) and *cDNAyft1.3* (4,793 bp) had the same 3,951 bp CDS and 260 bp 3′ UTR regions, but while the former had a 1,207 bp 5′ -UTR with a 560 bp In/Dels), *cDNAyft1.3* comprised a 582 bp 5′ UTR, from which most of the inserted DNA and part of u2.1 (5′ end of 123 bp at u2) in the 5′ UTR was absent (Supplemental Figure S4). The CDS derived from *cDNAyft1.3* was identical to that of *cDNA yft1.2* and the full length M82 *YFT1* cDNA. The expression of *cDNAyft1.1* was significantly higher than both *cDNAyft1.2* and *cDNAyft1.3* (Supplemental Figure S5), and we concluded that the *YFT1* allele undergoes alternative splicing to produce three different transcript types due to the 573 bp insertion event in the 5′ UTR (Supplemental Figure S4).

**Figure 8.**
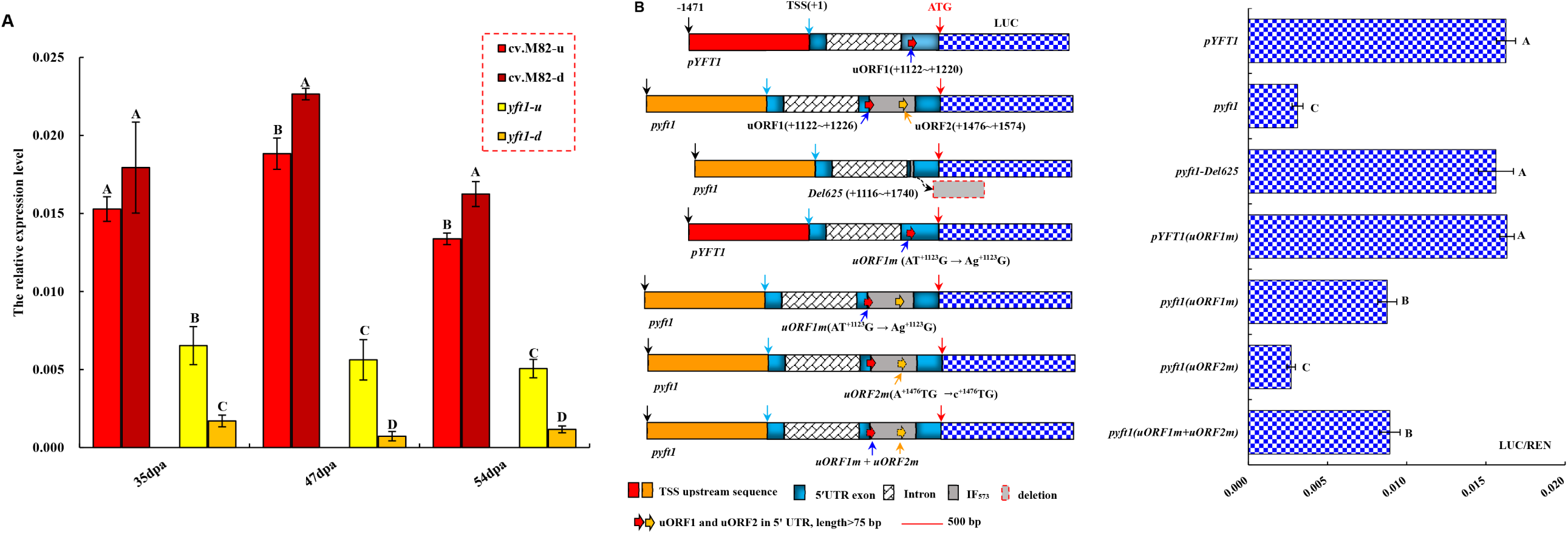
Insertion of the 573 bp DNA segment caused aberrant splicing and decreased *YFT1* allele transcripts in *yft1*. **(A**) Expression difference between up-/ downstream sequences of the broke site by *IF*_*573*_ in *YFT1/YFT1* allele of M82 / *yft1.* **M82**, wild type tomato. ***yft1***, *yellow-fruited tomato 1*(*n3122*) derived from M82 via fast-neutron irradiation. **M82-u/-d**, indicate the up-/downstream sequence of -318 bp (ATG start codon assigned to +1), determined by specific primers. ***yft1*-u/-d**, expression of up-/downstream sequence of -318 bp in *YFT1* allele in *yft1* using the same primers as for M82. **(B)** *uORF* affects promoter strength. **+1**, assigned to the first letter (adenine, A) of the transcription start site (TSS). ***pYFT1***, *YFT1* gene promoter, amplified from a DNA sequence 3,000 bp upstream of ATG in M82. ***pyft1***, *YFT1* allele promoter, a 3,560 bp DNA fragment upstream from ATG in *yft1*, containing two mutations: an insertion of 573 bp and a deletion of 13 bp. ***pyft1-Del625*** (+1116 to +1740), a 625 bp DNA fragment consisting of 84 bp (3’ end of *u2.1*) and 541 bp (5’ end of *IF*_*57*_***3***) removed from *pyft1.* The deleted DNA fragment corresponds to intron 2 in the *yft1.3* transcript. ***pYFT1*(*uORF1m*)**, changed AT^+1123^G into A*g*^+1123^G in *pYFT1* to mutate the ATG of uORF1(+1122 to +1220). ***pyft1*(*uORF1m*)**, converted AT^+1123^G into A*g*^+1123^G in *pyft1* to mutate ATG of uORF1(+1122 to +1226). ***pyft1*(*uORF2m*)**, converted A^+1476^TG into *c*^+1476^TG in *pyft1* to mutate uORF2(+1476 to +1574). ***pyft1* (*uORF1m*+*uORF2m*)**, *pyft1* harboring both the *uORF1m* and *uORF2m* mutation. All the mentioned ORFs in the figure were identified using an ORF finder (https://www.ncbi.nlm.nih.gov/orffinder/). **ATG**, translation initiation codon. ***LUC***, gene encoding luciferase. Data indicate the means ± SD (n=3). Capital letters indicate statistical significance at the *P* <0.01 level based on a Duncan’s test.

### *IF*_*573*_ breaks the 5’-UTR structure

A comparison of the genomic DNA structure and full size cDNAs of the *YFT1* allele in *yft1* with *YFT1* in M82 revealed a genetic lesion in the second upstream exon (uE2) of the *yft1* 5’-UTR. This was predicted to alter the first open-reading frame (uORF_1_) structure in the uE2 (Supplemental Figure S4). The stop codon (TSC) of the uORF acts as a premature termination codon (PTC) that causes nonsense-mediated mRNA decay (NMD) in the affected gene (Spellman et al., 2007, Ge and Porse, 2014). To identify alterations in the structural organization of uE2 in *yft1*, we used an ORF finder to identify uORFs with a length >75 bp in uE2. This resulted in identification of one *M82-uORF1* (99 bp) from M82 and *yft1-uORF1* (105 bp) and *yft1-uORF2* (99 bp) from *yft1*. We also found that the first 78 bp DNA sequence of *yft1-uORF1* was identical to that of *M82-uORF1*, while the *yft1-uORF2* was derived from *IF*_*573*_ in *yft1*.

To investigate whether the promoter strength was affected by variation in the uORF structures, a series of mutated promoters were created to disrupt the uORFs through mutation of the ATG start codon. This series consisted of: *pyft1-Del625, pyft1* (*uORF1m*), *pyft1* (*uORF2m*), *pyft1* (*uORF1m*+*uORF2m*), and *pYFT1*(*uORF1m*). A difference was not detected in the LUC/REN ratio between *pYFT1, pYFT1*(*uORF1m*), and *pyft1-Del625* in transiently transformed MicroTom (*S. lycopersicum*) fruit, as well as *pyft1* vs *pyft1*(*uORF2m)*, and *pyft1*(*uORF1m*) vs *pyft1*(*uORF1m*+*uORF2m*). However, the LUC/REN ratio in *pyft1*(*uORF1m*) /*pyft1*(*uORF1m*+*uORF2m*) containing *IF*_*573*_ was significantly lower than in *pYFT1, pYFT1*(*uORF1m*) and *pyft1-Del625*, and statistically higher than in *pyft1*/*pyft1*(*uORF2m).* This suggested that *yft1 uORF1* plays a key role in the downregulation of the *YFT1* allele in *yft1*, and likely acts as a PTC to induce NMD in the *YFT1* allele (Fig. 8B). In conclusion, the strength of the *YFT1* allele promoter in *yft1* was associated with both *IF*_*573*_ and the structure of the 5’-UTR (Fig. 8B).

## Discussion

The genetic lesion of *YFT1* in *yft1* was mapped to a region of 1,200 bp to 1,772 bp downstream of the transcription start site (TSS) (Supplemental Figure S4), and the candidate *EIN2* gene was confirmed here through genetic complementation (Fig. 1). Expression of the *YFT1* allele is seriously impaired in *yft1*, to 3% of M82 levels in stamens and ∼5% in fruit (Supplemental Figure S1). The genetic lesion resulted in 5,053 DEGs between *yft1* and M82 fruits (Fig. 3), as well as impairment of chromoplast development (Fig. 4), carotenoid accumulation (Fig. 5), and ethylene synthesis and signal transduction (Fig. 6).

To investigate the mechanisms underlying the downregulated *YFT1* expression, promoter analysis was performed, which revealed that the deletion of two functional regions in artificial promoters were responsible (Supplemental Figure S3). *IF*_*573*_ is present in the 5’-UTR of *yft1* and we associated the downregulated expression with three factors. Firstly, two negatively regulatory sequences were identified, and mapped to regions from 273 bp to 173 bp and 172 bp to 73 bp in *IF*_*573*_ (Fig. 7). Secondly, aberrant splicing sites were introduced into the *YFT1* 5’-UTR by the *IF*_*573*_ event, triggering alterative splicing events. Three types of cDNAs (*yft1.1, yft1.2*, and *yft1.3*) were detected in *yft1* (Supplemental Figure S4), and the abundance of *cDNA yft1.1*, which is not predicted to encode a functional protein, was significantly higher than that of both *cDNA yft1.2* and *yft1.3* in all organs of *yft1*, except leaf and fruit (Supplemental Figure S5). In contrast, only one type of *YFT1 cDNA* was identified in M82 (Supplemental Figure S4). The expression of *YFT1* in *yft1* was lower than in M82 (Supplemental Figure S1), suggesting that only a few YFT1/EIN2 proteins were synthesized in *yft1* compared with M82, which would impede transduction of the ethylene signal. Thirdly, the second 5’-UTR exon in *yft1* was disrupted due to the *IF*_*573*_ event, which increased the *uORF*_*1*_ length from 99 bp (from 1,122 bp to 1,220 bp downstream of the TSS in M82) to 105 bp (from 1,122 bp to 1,226 bp downstream of the TSS in *yft1*), resulting in a novel *uORF*_*2*_ (from 1,476 bp to 1,574 bp downstream of the TSS) in the *yft1* region. Although the length of *uORF*_*1*_ (105 bp) in *yft1* was different (99 bp) from that in M82, the DNA sequence of the first 78 bp was identical. However, a uORF_2_ of 99 bp derived from *IF*_*573*_ was only detected in *yft1*, and not in M82. Point mutations in the promoter showed that promoter strength was not associated with either *M82-uORF*_*1*_ or *yft1-uORF*_*2*_, while *yft1-uORF*_*1*_ played a key role in the downregulation of *YFT1* (Fig. 8B). Intron retention is often accompanied by NMD (Zhan, 2013; Ge and Porse, 2014), and a TSC sequence in the upstream uORF frequently acts as a PTC to cause NMD in the affected gene (Spellman et al., 2007). In this study, the TSC of uORF_1_ in both M82 and *yft1* was abolished by a point mutation in the ATG start codon, indicating that the downregulation of *YFT1* in *yft1* could be attributed to alteration of *uORF*_*1*_ due to the *IF*_*573*_ event. The uORFs likely trigger mRNA degradation in a size-dependent manner (Nyikó et al., 2009).

The YFT1/EIN2 protein is a central constituent in ethylene signal transduction, and genetic lesion of *YFT1* in *yft1* resulted in the differential expression of 5,053 genes compared with M82. These genes were annotated as being involved in physiology, biochemistry and development, and especially impaired ethylene synthesis and signal transduction, delayed chromoplast development, and decreased carotene accumulation, resulting in a yellow-fruited *yft1* phenotype.

## Conclusions

An insertion event of 573 bp at 1,200 bp downstream of the TSS in *YFT1* allele of *yft1* mutant tomato impaired its own transcriptional expression, and caused in yellow-fruited phenotype in *yft1* mutant. Our research revealed the mechanism behind the downregulated *YFT1* expression in *yft1.* These results contribute to the understanding of fruit color formation via regulation of *YFT1*/*EIN2*, which acts as a central node connecting processes such as ethylene synthesis and signal transduction, chromoplast development, and carotenoid accumulation.

## Materials and methods

### Plant materials and growth conditions

*yft1* (*n3122*) and wild type (WT; cv. M82) seeds were kindly provided by Prof. Dani Zamir (http://zamir.sgn.cornell.edu/mutateds). The resulting plants, as well as the overexpression and silenced *YFT1-CDS* transformed tomato lines, were grown under a standard greenhouse conditions at the Pujiang experimental base, Shanghai Jiao Tong University, Shanghai, China.

### Constructs and genetic transformation of tomato

The CDS (coding sequence) of *YFT1* was amplified from a red-fruit M82 cDNA library using the primers *YFT1-CDS-F* and *YFT1-CDS-R* (Supplementary Table S1), and the resulting *YFT1-CDS* fragment was then ligated into the *pHB* plasmid (Mao et al., 2005) at the *BamH* I and *Xba* I sites using the In-Fusion kit (Vazyme Biotech Co., Ltd, Nanjing, China), to create the binary expressional vector *2*×*35S::YFT1-CDS* (called *YFT1-CDS*). A 280 bp DNA segment was also amplified using the *RNAi-1549-F* and *RNAi-1836-R* primers (Supplementary Table S1) and cloned into the *pHELLSGATE12* plasmid using the Gateway system (Invitrogen, CA, USA) to generate *35S::YFT1-RNAi* (referred to as *YFT1-RNAi*). The *YFT1-CDS* and *YFT1-RNAi* constructs were transformed into *Agrobacterium tumefaciens* (strain EHA105) using the freeze-thaw method (Weigel and Glazebrook, 2006) and the engineered strains were used to transform *yft1* and M82 tomato lines, respectively.

*yft1* and M82 seeds were immersed in 75% ethanol (v:v=3:1) for 2 min, and then sterilized in 20% sodium hypochlorite (v:v=1:4) for 10 min at room temperature, before washing at least five times with sterile water for 5 min. The sterilized seeds were sown on MS_0_ medium containing 4.41 g/L MS powder (Sigma, CA, USA), 30 g/L sucrose, 2.6 g/L phytagel (Sigma, CA, USA) (pH 5.8) and germinated with a light period of 16 h/8 h (light/ dark) at 25°C.

Tomato seedlings (cotyledon or hypocotyl) were cut into ∼0.3 cm explants (leaf disks or cotyledon extensions), and pre-cultivated on MS_1_ medium[MS_0_ with 0.1 g/L IAA (Sangon Biotech Co., Ltd, Shanghai, China) and 1.5 g/L 6-BA (Sigma, USA)], under dark conditions at 25 °C for 24 h. The EHA105 strain harboring *YFT1-CDS* and *YFT1-RNAi* were cultured to OD_600_=0.6 in lysogeny broth (LB) medium at 28 °C with gentle (250 rpm) shaking, and re-suspended in an equal volume of MS_0_ medium without phytagel at 28°C, 180 rpm for 30 min, before addition of acetosyringone (Sangon Biotech Co., Ltd, Shanghai, China) to final concertation of 50 mg/L. Tomato explants were transferred to a glass flask with 50 mL of the *Agrobacterium* suspension and gently shaken for 6 min. The explants were then co-cultured on MS_1_ medium at 25°C for 36 h under dark conditions after removal of *Agrobacterium*. The explants were then transferred to MS_2_ medium, consisting of MS_1_ with 300 mg/L carbenicillin and 5 mg/L hygromycin (Roche, Germany) (*YFT1-CDS*), or 50 mg/L kanamycin (*YFT1-RNAi*). The MS_2_ medium was replaced with fresh medium at 12 to 14 days intervals. The regenerated seedlings were cut from the explants to root on MS_3_ medium (MS_0_ with 300 mg/L carbenicillin and 5 mg/L hygromycin for *YFT1-CDS* or 30 mg/L kanamycin for *YFT1-RNAi*) when they reached 3∼5 cm. Finally, the rooted seedlings were transferred into a pot with soil to acclimatize in a plant culture chamber with an 8 h/16 h dark/light photoperiod at 25 °C for 10 days, and then grown under standard greenhouse conditions.

### Transcriptome analysis

Total RNA was extracted from *yft1* and M82 pericarp from fruit that were harvested at 35 days post anthesis (dpa; Mature Green stage, MG), 47 dpa(Breaker stage, BR) and 54 dpa(Red Ripe stage, RR) using the RNAprep pure Plant Kit (Tiangen, Beijing, China) based on the manufacturer’s instructions. Total RNA samples were used to construct cDNA libraries for paired-end RNA sequencing (2×100 base pairs) (Wang et al., 2014) on an Illumina HiSeq 2000 platform by the Shanghai Majorbio Bio-pharm Biotechnology Co. (Shanghai, China).

The FASTX-Toolkit (http://hannonlab.cshl.edu/fastx_toolkit) was used to analyze the raw reads and remove the adaptor sequences and low-quality regions. The remaining high-quality reads were aligned to the tomato reference genome SL2.40 (ITAG2.3, ftp://ftp.solgenomics.net/tomato_genome/annotation/ITAG2.3_release/) using TopHat (version 2.0) (Trapnell et al., 2009). The RNA abundance for each gene was then evaluated using the FPKM (fragments per kilo-base of exon per million fragments mapped) determined by Cufflinks (http://cufflinks.cbcb.umd.edu/) (Trapnell et al., 2010). Genes with FPKM values ≥1 in at least one sample were used to identify the differentially expressed genes (DEGs) with a log2 (Fold Change) cutoff value > 1 and a FDR (false discovery rate) < 0.05.

GO (Gene Ontology) classification and analysis were carried out using Blast2GO (Conesa et al., 2005), and the significantly enriched GO terms amongst the DEGs were identified by Fisher’s exact test with FDR <0.05. The DEGs were used in BLAST searches of the KEGG database (http://www.genome.ad.jp/kegg/kegg2) to annotate pathways with E-value ≤ 10^−50^ (Altschul et al., 1997). The KEGG pathway enrichment analysis was also subjected to a Fisher’s exact test with FDR < 0.05.

### Real time –quantitative PCR (RT-qPCR) analysis

Total RNA (1 μg) extracted from pericarp of fruit of the genotypes M82, *yft1, YFT1-CDS*, and *YFT1-RNAi*, at 35 dpa, 47 dpa and 54 dpa, was used to generate templates to synthesize first chain cDNA using the PrimeScript™ RT Master Mix kit (TAKARA, Dalian, China), with three biological replicates sample. The cDNA was diluted 50 times with double distilled water to provide the template for the RT-qPCR analysis. Twenty μL reaction volumes containing 10 μL SYBR premix ExTaq (TAKARA, Dalian, China), 0.6 μL of each of the forward and reverse primers, 2 μL cDNA template, and 6.8 μL ddH_2_O were used and the RT-qPCR was performed using a Light Cycler 96 Real-Time PCR Machine (Roche, Germany). Expression levels were calculated using the 2^-ΔCT^ equation (Kilambi et al., 2013) with *ACTIN* (GenBank accession: BT013524) as an internal reference gene for normalization. The amplification program was as follows: 2 min for initial denaturation at 94°C; 40 cycles for 20 s at 94°C, 20 s at 55°C, and 20 s at 72°C. Gene specific primers (forward/reverse) are listed in Supplementary Table S1.

### Measurement of carotenoid content

Pericarp from the equatorial region of the fruit was sampled at 35 dpa, 47 dpa and 54 dpa in three biological replicates, cut into small pieces and powdered in liquid nitrogen using a mortar and pestle. Carotenoids were extracted using the protocol described in Fraser *et al*. (2000), with minor modifications. Approximately 0.5 g of powdered pericarp was homogenized in a solution of 1.5 mL KOH and methyl alcohol (w/v = 6%) and incubated at 60 °C for 30 min. An aliquot of 1.5 mL Tris buffer (50 mM Tris-HCl, 1 M NaCl, pH 7.5) was added and the solution was gently mixed by inverting the tube, before incubation at 4 °C for 10 min and addition of 4 mL chloroform. This solution was also mixed by inversion and incubated in an ice bath for 10 min, before centrifugation at 3,000 g at 4 °C for 10 min. The samples were extracted twice with chloroform and the lower organic phase was collected, dried in a SPD 2010 vacuum concentrator (Thermo Fisher Scientific Co., Ltd., MA, USA), and the samples re-dissolved in 50 μL methyl tertiary butyl ether (MTBE) (Thermo Fisher Scientific, MA, USA), of which 1.0 μL was injected into a Waters Acquity Ultra-performance Convergence Chromatography (UPC^2^) system (Waters Corp., Milford, MA, USA) to determine carotenoid content. The detection parameters were as described in Li et al. (2015), with minor modifications. The temperature in the sample chamber was kept at 10 °C, the Waters ACQUITY UPC2 HSS C18 SB (100 mm×3.0 mm, 1.8 μm) was used for carotenoid separation. The binary mobile phase was composed of CO_2_ (A) and methanol/ethanol (v/v=1:2) (B). The linear elution gradient was 0.5 min, 95% A + 5% B; 2 min, 70% A + 30% B; 5 min, 70% A + 30% B; 5.5 min, 95% A + 5% B; 7 min, 95% A + 5% B. A flow rate of 1.5 mL/min was used and the column temperature was set to 45°C with a backpressure of 22.9 MPa. The PDA (photo-diode array) detector operated from 210 nm to 500 nm with wave length compensation from 210 nm to 280 nm. The Empower software (version 3, Waters Corp., Milford, MA, USA) was used for instrument control and data acquisition.

Lycopene, β-carotene and α-carotene standards were purchased from the Sigma Chemical Co. (CA, USA) and lutein from the Yuanye Biotechnology Co., Ltd. (Shanghai, China). The carotenoid standards were prepared as follows: α-carotene, β-carotene, lutein and lycopene were dissolved in MTBE at a 500 μg/mL concentration. Standard dilution series (5 μg/mL, 10 μg/mL, 50 μg/mL, 250 μg/mL, 500 μg/mL) were made to establish standard curves. Carotenoid concentrations in the extracts were determined by comparison with the standard curves.

### Visualization of chromoplast ultrastructure using transmission electron microscopy (TEM)

Pericarp was sampled from M82 and *yft1* and the transgenic *YFT1-CDS* and *YFT1-RNAi* lines at 35 dpa, 47 dpa, and 54 dpa in three biological replicates, and the chromoplast ultrastructure was visualized using TEM (Tecnai G2 Spirit Biotwin, USA). Samples were prepared as described in Schweiggert et al. (2011) with minor modifications. The tomato pericarp was cut into 1 mm^3^ pieces using a scalpel and immediately immersed in 2.5% glutaraldehyde (v: v: v=25% glutaraldehyde: 0.1 M phosphate buffer: ddH_2_O=1:5:4, pH 7.3∼7.4) and then vacuumed infiltrated using a vacuum drying cabinet (Yiheng, Shanghai, China) with a vacuum pump (Millipore, MA, USA) for 30 min, before fixing at 4 °C for 6 h. The pericarp samples were then washed three times with 0.1 M phosphate buffer (pH 7.2) for 15 min.

The pericarp samples were immersed in 0.1 M phosphate buffer (pH 7.2) containing 2% osmium tetroxide (w:v=2%) for 2 h, and washed three times 15 min using 0.1 M phosphate buffer (pH 7.2). The samples were then dehydrated using gradient series of ethanol (50%, 70%, 90% ethanol) with a mixture of 90% acetone and 90% ethanol (v:v=1:1) wash, with each incubation lasting for 15 min. Finally, the samples were transferred into 100% acetone and shaken at 100 rpm (Qilinbeier Instrument Manufacturing Co., Ltd, Jiangsu, China) at room temperature for 20 min. This was repeated twice with fresh acetone. After dehydration, the pericarp samples were immersed in a mixture of acetone and epoxy resin 812 (Hede Biotechnology Co., Ltd., Beijing, China) (v:v=1: 1) for 1 h and overnight (v:v=1: 2), before a final immersion in 100% epoxy resin 812 for 7 h, and transferred to embedding plates for 48 h at 60°C. Sections (70 nm thick) were generated using a Leica UC6-FC6 microtome with a diamond knife (Leica Microsystems Inc., Germany), and stained with 2% lead citrate (w:v=2%) (Zhongjingkeyi Technology Co., Ltd., Beijing, China), before imaging using a 120 kV biological TEM (FEI, OR, USA).

### Ethylene emission

Tomato fruit were collected from the *YFT1-CDS, YFT1-RNAi*, M82 and *yft1* lines at three development stages (35 dpa, 47 dpa, and 54 dpa) in three biological replicates. After weighing, the tomato fruit were kept at 25 °C for 2 h in a thermostatic incubator (Tensuc Lab Instruments Manufacturing Co., Ltd., Shanghai, China) to reduce the effects of harvesting stress-induced ethylene synthesis. Subsequently, the fruit were sealed in an airtight 500 mL bottle at 25 °C for 4 h and 1 mL gas samples were removed by an injector and injected into a GC-2010 gas chromatograph (Shimadzu Co., Ltd., Japan). The instrument parameters were set as follows: temperature of injector and flame ionization detector set at 200 °C and 250 °C, respectively; initial temperature set to 40 °C for 5 min and increased to 220 °C at a rate of 30 °C/min. The endogenous ethylene concentration (nL·g^-1^·h^-1^) was calculated using an ethylene standard curve established based on the peak area derived from six different volumes (0 mL, 0.05 mL, 0.1 mL, 0.3 mL, 0.5 mL, 0.6 mL) of 10 μL·L^-1^ ethylene (Weichuang standard gas analytical technology Co., Ltd., Shanghai, China).

### Response to ethylene

Sterilized *YFT1-CDS, YFT1-RNAi*, M82 and *yft1* tomato seeds were sown separately on MS_0_ solid medium containing 4.41 g/L MS powder (Sigma, CA, USA), 30.00 g/L sucrose, 2.60 g/L phytagel (pH 5.8) with/without 10 μM 1-aminocyclopropane-1-carboxylic acid (ACC) (Sigma, CA, USA), and grown under dark conditions at 25 °C. After 7 days, the lengths of the hypocotyls and roots were measured using CD-15CPX Vernier caliper (Mitutoyo, Japan), and photographed using an EOS 800D single lens reflex (SLR) digital camera (Canon, Japan).

### Sequencing of the full-length *YFT1* cDNA

Total RNA samples extracted from M82 and *yft1* pericarp (47 dpa, BR stage) using the RNAprep pure Plant Kit (Tiangen, Beijing, China) were used as a templates in rapid amplification of cDNA ends (RACE) analyses, according to the manufacturer’s instructions (SMARTer® RACE 5’/3’ Kit, Clontech, USA), with gene specific primers (*YFT1*-GSP5′-R/GSP3′-F, Nest-F(5’)/R(3’) and GSP_573_-R(5’)/F(3’)) were designed and synthesized by Sangon Biotech (Shanghai, China) (Supplementary Table S1). The PCR products were separated on a 1.0% agarose gel in 1×Tris-Acetate-EDTA (TAE) buffer, and the gel area containing the target DNA band was purified using the Axygen DNA gel extraction kit (Axygen, China), before ligation into the pRACE vector for sequencing.

### Detection of chimeric *YFT1* promoter activity

Based on the 3 kb upstream sequence from the predicted start codon ATG in *YFT1* (M82), primers were designed and synthesized (*pYFT1-Sal* I*-F* / *pYFT1-Nco*I*-R*) to amplify the *pYFT1* promoter sequences from M82 and *pyft1* (Supplementary Table S1). In addition, three mutated promoters of *pYFT1-Del13, pfyt1*+*Del13u* and *pfyt1*+*Del13d* were created using the designed overlap primers (Supplementary Table S1) by overlap PCR (Higuchi et al., 1988). These promoter fragments were separately ligated into at sites of *Sal* I and *Noc* I in the *pCAMBIA1391* plasmid (stored in Zhao Lab, Shanghai Jiao tong University, China), upstream from the β-glucosidase (*GUS*) gene.

Subsequently, a series of chimeric promoters was created based on the CaMV 35S promoter fused with different lengths of DNA sequences derived from the 573 bp insertion segment (called *IF*_*573*_). These were named *35S-IF*_*573*_, *35S-IF*_*473*_, *35S-IF*_*373*_, *35S-IF*_*273*_, *35S-IF*_*173*_, and *35S-IF*_*73*_ with the numbers indicating how many base pairs of the 3’ terminal end of *IF*_*573*_ were included. *35S* and *IF*_*573*_ served as positive and negative control promoters, respectively. These chimeric promoters were all fused to upstream of the *Firefly Luciferase* gene using *Sal* I and *Noc* I sites in the pGreen II 0800-LUC plasmid (Hellens et al., 2005).

The constructs were introduced into *Agrobacterium* strain GV3101 using the freeze-thaw method (Weigel and Glazebrook, 2006), and cultured to an OD_600_ value of 0.6 by shaking at 250 rpm and 28 °C. The cells were collected by centrifugation at 3,000 g and re-suspended in 1/2 MS_0_ medium containing 2.21 g MS powder and 30.00 g sucrose per liter, with acetosyringone and 2-(N-Morpholino) ethanesulfonic acid (MES) to a final concentration of 40 μM and 10 mM. The re-suspended cells were used to transform the leaves of 6 weeks old tobacco (*Nicotiana benthamiana*) seedlings grown in a chamber at 25 °C with a light period of 16 h/8 h (light/dark) by culture infiltration using a needleless syringe. The transformed tobacco plants were subsequently transferred to a chamber and grown at a photoperiod of light/dark (24 h/24 h) at 25 °C. Disks (0.3 cm diameter) were punched from the transformed regions of the leaves and powdered in Eppendorf tubes using liquid nitrogen and two steel balls with a Tissue Lyser (WanBai biotechnology Co., Ltd., Shanghai, China). Measurement of GUS activity was as described in Jefferson (1987) and luciferase activity was detected using a GloMax (Promega, WI, USA) Luminometer, according to the manufacturer’s protocol.

### Dual-luciferase activity assay

To test the promoter activity, a series of mutated promoters was created by deletion or point mutation of the ATG start codon (over 75 bp) in the 5’-UTR of both *pYFT1* and *pyft1*. There were *pyft1-Del625* (with deleted 625 bp DNA segment from 1,200 bp to 1,740 bp downstream of the transcription start site in *pyft1*), as well as *pYFT1*(*uORF1m*), *pyft1*(*uORF1m*), *pyft1*(*uORF2m*), and *pyft1*(*uORF1m*+*uORF2m*), where *uORF1m* and *uORF2m* indicate changing AT^**+1123**^G to A*g*^**+1123**^G, and A^**+1476**^TG to *c*^**+1476**^TG, respectively. The promoter segments were inserted into the pGreen II 0800-LUC plasmid using the *Sal* I and *Nco* I sites located upstream from the *Firefly* and *Renilla* genes. The constructs were introduced into *Agrobacterium* (strain GV3101) and used to transform fruit of the MicroTom (*S. lycopersicum*) genotype. *pYFT1* and *pyft1* served as positive and negative control promoters, respectively.

GV3101 harboring pGreen II 0800-LUC was cultured to an OD_600_ culture density of 1.0 in liquid LB medium with 100 mg/L kanamycin (Kan), 50 mg/L rifampicin (Rif)and 30 mg/L gentamicin (Gen), shaking at 250 rpm at 28 °C. The cells were transferred to an induction medium [of 0.5% beef extract (Sangon Biotech Co., Ltd, Shanghai, China), 0.1% yeast extract, 0.5% peptone, 0.5% sucrose, 2 mM MgSO_4_, 20 mM acetosyringone, and 10 mM MES, pH 5.6] and then cultured to an OD_600_ of 1.0. Cells were collected by centrifugation at 3000 g for 15 min and equal volume re-suspended in infiltration medium (10 mM MgCl_2_, 10mM MES, 200 mM acetosyringone, pH 5.6), before incubation on a THZ-C shaker (Taicang Experimental Equipment FTY., Jiangsu, China) at 20 rpm at room temperature for 2 h. Transient transformation of tomato fruit was performed as described in Orzaez et al. (2006), with minor modifications. Briefly, infiltration was conducted in three biological replicates using a 1 mL syringe. The needle was inserted 3∼4 mm from the stylar apex and the *Agrobacterium* culture was gently injected until the infiltration solution appeared at the tip of the sepals. The inoculated fruits were sampled excluding seeds and damaged parts after 24 h in the dark or and 48 h in the light at 25 °C, and powdered in liquid nitrogen. Luciferase activity was detected using a GloMax (Promega, WI, USA) Luminometer according to manufacturer’s protocol (Promega, WI, USA).

### Accession Numbers

The RNA-seq data analysed in this study were deposited in the Gene Expression Omnibus (GEO) database, http://www.ncbi.nlm.nih.gov/geo (Accession no. GSE73908)

## Supplemental Data

The following materials are available.

**Supplemental Table 1.** Primers used in the present study.

**Supplemental Figure S1.** *YFT1* expression patterns in M82 and *yft1.*

**Supplemental Figure S2.** Position of an inserted DNA fragment of 573 bp (*IF*_*573*_) at chromosome 6.

**Supplemental Figure S3.** Identification of the causal event leading to downregulated expression of *YFT1* in *yft1.*

**Supplemental Figure S4.** Diagram of the *YFT1/YFT1* allele in M82*/ yft1.*

**Supplemental Figure S5.** Expressional difference among transcripts of three types derived from *YFT1* allele in *yft1*.

## Acknowledgements

The authors are grateful to Dr. Dani Zamir and the TGRC (Tomato Genetic Resource Center, UC-Davis, CA, USA) for providing tomato seeds. We thank Dr. Xin Li and Ge Wang (Instrumental Analysis Center, Shanghai Jiao Tong University, Shanghai, China) for help with the carotenoid analysis and TEM analysis.

